# Mining the Proteome of Human Ovarian Cancer Extracellular Vesicles Using Thermolysin Proteolysis

**DOI:** 10.1101/2024.10.10.617562

**Authors:** Tyler T. Cooper, Jaihui Lui, Olena Bilyk, Michael Jewer, AOCS Study Group, Helen Steed, Yangxin Fu, Lynne-Marie Postovit, Gilles A. Lajoie

**Author notes:** Co-senior Authors (Lynne-Marie Postovit, Gilles A. Lajoie).

## Abstract

This study investigates the utility of Thermolysin as a proteolytic enzyme to enhance the depth and coverage of proteomic analysis in ovarian cancer (OC) extracellular vesicles (EVs). EVs were isolated from OC cell lines and ascites fluid samples from women diagnosed with high-grade serous carcinoma. Proteins were digested using Thermolysin and Trypsin/LysC, followed by label-free data-dependent acquisition (DDA) and data-independent acquisition (DIA) mass spectrometry. The proteolytic efficiency, sequence coverage, peptide complexity, and proteomic depth were compared between Thermolysin and Trypsin/LysC digests. In silico analyses predicted theoretical benchmarks of these parameters using a core set of 22 proteins, and Gene set enrichment analyses (GSEA) highlighted the biological relevance of proteins identified throughout the study. Thermolysin digestion significantly increased the complexity or of peptide pools compared to Trypsin/LysC leading to limited peptide and protein identification, albeit total sequence coverage was increased through complementation to tryptic peptides. In both cell line and ascites EVs, Thermolysin identified unique proteins not detected by Trypsin/LysC that are known drivers of metastatic solid cancers, such as Ly6E. Offline strong cation exchange (SCX) fractionation improved proteomic depth and sequence coverage obtained with Thermolysin to generate a spectral library for DIA. Our DIA analysis of Thermolysin digests revealed the presence of NODAL in OC ascites EVs, a protein associated with poor clinical prognosis, which was not detected in Trypsin/LysC digests. The importance of NODAL was cross-validated in TGCA-OV and AOCS datasets by clinical cohorts by assessing RNA levels in solid tumors or ascites fluid, respectively. Collectively, we demonstrate that Thermolysin complements traditional enzymes like Trypsin/LysC to provide a more comprehensive proteomic landscape for biomarker discovery.

## Introduction

Tandem mass spectrometry (LC-MS/MS) is a powerful tool for the characterization and quantification of proteins in complex biological samples^1^. As of this writing, the sequencing of the human genome is complete, with an estimated 13,506 protein-coding regions. However, the human proteome remains less resolved, despite the presence of over 20,000 unique proteins listed in the human UniProt database (20,465 entries as of January 1st, 2024). The complexity of the human proteome is further amplified by post-translational modifications^2^ (e.g., phosphorylation) and proteoforms generated through alternative splicing of mRNA^3^ or cleavage from pro- to active forms^4^. Most often, a “bottom-up” approach is used in LC-MS/MS to identify and quantify thousands of proteins in a single “shotgun” experiment through the detection of ionized peptides in either an orbitrap^5^ or linear ion trap mass analyzer^6^. Shotgun analysis of label-free and isobaric-tagged peptides does not require the complete protein sequence to be identified for confident protein annotation and quantification^7^. While this feature has led to thousands of biological discoveries, it impedes the capacity to fully ‘mine’ the proteome of complex biological samples^8, 9^. For instance, in-solution digestion of HeLa proteomes with Trypsin/LysC achieves only about 10% sequence coverage per protein, which can be enhanced to ∼60% by combining MS/MS spectra from multiple offline fractionation techniques^10^. The caveat of offline fractionation is the increase sample preparation time and increased mass spectrometry time even with the addition of high-cost isobaric labelling. In addition to offline fractionation of peptides prior to LC-MS/MS, the development of new MS^n^ technologies^11–13^, online fractionation by ion mobility^14, 15^, and alternative fragmentation techniques has been proposed to increase sequence coverage across the whole proteome^16^. Despite these advances, less than half of the peptides produced by trypsin are suitable for LC-MS/MS due to biophysical properties or poor ionization^17^. Using alternative enzymes to Trypsin can also increase sequence coverage by detecting adjacent or inaccessible peptide sequences^16^, warranting the investigation of uncommon enzymes to enhance coverage during bottom-up or even middle-down LC-MS/MS.

Proteins localized at the plasma membrane of cells often contain cytoplasmic, extracellular, and transmembrane domains, with topography determined by local amino acid characteristics, such as hydrophobicity. Unfortunately, Trypsin often does not provide optimal sequence coverage for most of these proteins, despite the presence of arginine and lysine. An estimated 75% of amino acids in transmembrane domains are comprised of hydrophobic residues, such as glycine (G), alanine (A), valine (V), leucine (L), isoleucine (I), proline (P), phenylalanine (F), methionine (M), and tryptophan (T)^18^. Thermolysin is a metalloproteinase, isolated from the Gram-positive bacteria *Bacillus thermoproteolyticu*, that targets the N-terminal of aliphatic amino acids (F/M/V/A/I/L) and functions at high temperatures (65-85°C) where secondary structures in proteins may be disrupted^19–23^. The number of proteomic studies employing Thermolysin are extremely limited, and none within the context of cancer research.

Extracellular vesicles (EVs) are secreted by almost all cell types and consist of a luminal core of biological cargo enclosed by a lipid bilayer derived from the plasma membrane^24, 25^. As a result, the proteomic cargo of EVs is contained within both the lumen and as transmembrane proteins, which may provide unique biological activity and serve as clinically relevant biomarkers for cancer detection^26^. Indeed, EVs may act a drug delivery systems^27^ or engineered for applications of regenerative medicine^28^. EV contain a diverse range lipid, nucleic acids and proteomic cargo^29, 30^, but most EV proteome studies do not focus on sequence coverage, post-translational modifications, or proteoforms. These features will be important to uncover as we aim to fully understand the therapeutic or diagnostic potential of EVs. We have previously isolated cell culture and biofluid EVs for prospective biomarker discovery using LC-MS/MS in parallel to surface enhance Raman spectroscopy^26, 30, 31^.

In our experience and aligned with others, Trypsin proteolysis alone is insufficient to provide complete sequence coverage for common EV proteins. For example, Trypsin cannot cleave within the transmembrane regions of CD9, limiting peptide matching to topographical domains. To address these limitations, we speculated that Thermolysin might provide sequence complementation to Trypsin by targeting hydrophobic amino acids within EV proteomes. To our knowledge, this is the first study to investigate Thermolysin as a proteolytic enzyme to enhance our capacity to ‘mine’ the EV proteome by increasing proteomic depth and complementary peptide identification for the aim of biological discovery. Herein, we highlight the benefits and speculate caveats of Thermolysin proteolysis for global proteomics.

## Material and Methods

### Cell Culture and Generation of Conditioned Media

OV-90 (ATCC® CRL-11732) and NIH:OVCAR3 (ATCC® HTB-161) were obtained from the ATCC. Human immortalized surface epithelial cells hIOSE (OSE364) were obtained from the Canadian Ovarian Tissue Bank at the BC Cancer Agency and kindly provided by Dr. Ronny Drapkin (Department of Obstetrics and Gynecology, University of Pennsylvania). Primary cell lines EOC6 and EOC18 were isolated from the ascites of patients with high-grade and low-grade serous ovarian cancer, respectively. All cell lines, except OVCAR3, were maintained in M199+MCDB105 supplemented with 5-15% FBS. NIH:OVCAR3 cells were cultured in RPMI-1640 supplemented with 20% FBS and 5µg/mL insulin. Media was exchanged with serum-free media for 20-30 hours to generate conditioned media (CM) for EV purification. All work involving patient samples (cell lines and ascites) was approved by the Health Research Ethics Board of Alberta-Cancer.

### High Grade Serous Carcinoma Ascites

All work involving patient samples (cell lines and ascites) was approved by the Health Research Ethics Board of Alberta-Cancer.

### Ultracentrifugation of Extracellular Vesicles

OC CM and ascites samples were first centrifuged at 200-300 x g at 4°C to pellet cells. Supernatants were diluted 1:10 in PBS (except CM) and centrifuged at 3,000 x g for 20 minutes at 4°C to remove cell debris. Large cellular fragments were pelleted from supernatants by centrifugation at 10,000 x g for an additional 20 minutes at 4°C and collecting the resulting supernatant. Finally, supernatants were ultracentrifuged at 120,000 to 140,000 x g (SW-28 rotor) for 2 hours at 4°C to pellet EVs on an OptimaTM L-100 XP ultracentrifuge (Beckman Coulter). The supernatant was removed, and EVs were resuspended in 100-300µL of PBS and stored at −80°C until further use. EVs analyzed in this study were utilized in parallel studies^26, 30^ and characterized according to ISEV guidelines^24^.

### Protein Precipitation and Digestion

To prepare CM for LC-MS/MS, ∼30µg of EVs were lyophilized to dryness and reconstituted in 8M Urea, 50mM ammonium bicarbonate (ABC), 10mM dithiothreitol (DTT), 2% SDS lysis buffer. Proteins were sonicated with a probe sonicator (10 × 0.5s pulses; Level 1) (Fisher Scientific, Waltham, MA), reduced in 10mM DTT for 30 minutes at room temperature (RT), alkylated in 100mM iodoacetamide for 30 minutes at RT in the dark, and precipitated in chloroform/methanol. Trypsin/LysC proteolysis was performed in 100µL 50mM ABC (pH 8) by adding Trypsin/LysC (Promega, 1:50 ratio) to precipitated EV proteins and incubated at 37°C overnight (∼20h) in a ThermoMixer C (Eppendorf) at 900 rpm before acidifying to pH 3-4 with 10% FA. Alternatively, Thermolysin proteolysis was performed in 100uL of 50mM ABC/10mM KCl (pH=8) by adding Thermolysin (Promega, 1:50 ratio) to proteins and incubating at 70°C for 1 hour at 900 rpm prior to acidification to 1% FA. Salts and detergents were removed from peptide samples using C18 stagetips made in-house. Briefly, 10 layers were stacked into 200µL pipette tips and rinsed with ice-cold methanol. Stagetips were conditioned with Solution A (80/20/0.1%; acetonitrile (ACN)/water/TFA), followed by Solution B (5/95/0.1%; ACN/water/TFA) prior to loading ∼20µg of peptides resuspended in Solution B. Duplicate washes were performed with Solution B prior to elution of peptides using Solution C (80/20/0.1%; ACN/water/FA) and final elution using a 50/50 mixture of ACN/0.1% FA. Peptides were centrifuged at 45°C under vacuum and resuspended in 0.1% FA prior to quantification by BCA and injection into LC-MS/MS.

### Offline Stagetip Strong Cation Exchange (SCX) Fractionation

25µg of peptides were loaded onto a 7-layer SCX stagetip, generated in-house using a 200µL pipette tip. Peptides were resuspended in 300mM NH4CH3CO2 (Ammonium Acetate). Briefly, stagetips were activated with methanol, washed with 0.1% TFA, and equilibrated with 300mM NH4CH3CO2. Bound peptides were loaded onto stagetips twice and washed with 0.1% TFA. Peptides from cell line EVs were eluted into four fractions: 50mM/5%ACN, 100mM/5%ACN, 250mM/5%ACN, and 5% NH4OH. Peptides from ascites EVs were eluted into six fractions: 75mM, 125mM, 150mM, 250mM, 300mM, and 5% NH4OH. Elutions from SCX stagetips were dried under vacuum, resuspended in water, dried again before resuspension in 0.1% FA for LC-MS/MS analysis.

### Data-dependent (DDA) UPLC-MS/MS

Digested peptides were resuspended in 0.1% formic acid prior to label-free LC/MS-MS. Approximately 1µg of each sample was injected onto a Waters M-Class nanoAcquity HPLC system (Waters, Milford, Massachusetts) coupled to a Q Exactive Plus Orbitrap (Thermo Fisher Scientific, Waltham, Massachusetts) operating in positive mode. Full LC-MS/MS parameters are outlined in Supplementary Tables 1. Data analysis was performed using PEAKS v11.0. Enzyme cleavage specificity to was set to specific, missed cleavages = 4, MS1 error = 20ppm, MS2 tolerance = 0.05Da, Carbamidomethylation as a fixed modification, Oxidation (M) and Acetylation (K, N-terminal) were set as variable modifications, and both peptide and proteins identification was set to an FDR = 0.01.

### Data-independent (DIA) UPLC-MS/MS

Using the same chromatography gradients as DDA analyses, 1µg of peptide was analyzed on QE Plus using wide-window data-independent analysis to survey peptides between 250-1000m/z. Average peak widths were determined to be 20 seconds; thus, we aim for a cycle time of 3 seconds to obtain ∼6 points across the curve. MS1 was obtained at a resolution of 60k (25 ms IT) followed by 50 windows of equal width (20m/z). The total cycle time was 2.7 ms. Raw MS files were processed in MSConvert using settings: PeakSelection=1 and ZeroSamples=-1. Peptides from DIA analyses were identified independent and dependent on the DDA-library produced from SCX-fractionated peptides. Data analysis was performed in PEAKS v11.0. Full LC-MS/MS Parameters are outlined in Supplementary Table 2.

### Clinical Validation Samples

OC datasets from TCGA were analyzed to create Kaplan–Meier (KM) curves of overall and progression-free survival, stratified by NODAL expression. Analysis was restricted to serous OC subtype regardless of TP53 mutation status, chemotherapy, and debulking surgery treatment groups. Sixty RNA samples from ascites of high-grade serous ovarian cancer (HGSOC) patients were received from the Australian Ovarian Cancer Study (AOCS). Detailed clinical information of AOCS patients is provided in Supplemental Table 3.

### Droplet Digital PCR (ddPCR) Assay

ddPCR (Bio-Rad) for detecting NODAL expression was conducted using TaqMan primer probe assay (Hs00415443_m1, Applied Biosystems). The number of target molecules detected was calculated using Quantasoft (Bio-Rad). For every sample, ddPCR was also used for detection of the housekeeping gene RPLP0 (4333761, Applied Biosystem).

### Data analysis

Proteomic data analysis and visualization was performed in a Python 3+ environment with supporting statistical and graphing libraries. Comparison to Vesiclepedia databases were performed using FunRich (version 3.1.3) software. Differential analysis was performed by either Mann-Whitney U test or Kruskal-Wallis H Test with Dunn’s post hoc test.

## Results

### In silico proteolysis predicts improved sequence coverage of common EV proteins

As this study is the first to mine the proteome of EVs using Thermolysin, we sought to establish a theoretical benchmark of EV proteins to assess the efficiency of proteome depth obtained during experimental analysis. The diversity of proteomic cargo in EVs is cell and context-dependent, so we focused *in silico* analysis on 22 proteins identified by Kugeratski *et al*. as ‘core’ EV proteins^32^ **(Figure 1A)**. Computational proteolysis allowed five missed cleavages to occur, as we found complete digestion with Thermolysin primarily generated small peptides irrelevant to protein identification (< 6 a.a. and/or z = 1). Complete digestion (0 missed cleavages) produced a similar number of peptides between Thermolysin and TrypLysC; however, an exponential divergence in peptide complexity was observed with increased missed cleavages. **(Figure 1B)**. Even at two missed cleavages, Thermolysin produced a two-fold increase in peptide complexity relative to TrypLysC as a result of generating more peptides per protein **(Figure 1C)**. Allowing five missed cleavages to occur generated on average ∼250 peptides per protein for TrypLysC, whereas Thermolysin generated ∼500 peptides per protein. While we anticipated missed cleavages generate larger peptides, it was surprising that TrypLysC generated larger peptides than Thermolysin **(Figure 1D)**. At maximum missed cleavages, peptide length was ∼2-fold longer in TrypLysC peptide pools compared to Thermolysin (30 vs 15 a.a.). The diversity of peptide charge (z) in Thermolysin samples increased with missed cleavages; however, average peptide charge remained comparable to TrypLysC **(Figure 1E)**. The theoretical hydrophobicity of peptides can be calculated using GRAVY scores, where positive scores indicate more hydrophobic peptides. Our *in silico* analyses indicated a positive correlation between Thermolysin with a high missed cleavage rate and hydrophobic GRAVY scores **(Figure 1F)**. On the other hand, peptides generated by TrypLysC became more hydrophilic with increasing missed cleavages. Using core EV proteins as a benchmark, complete sequence coverage was obtained in all proteins when at least four missed cleavages were induced in Thermolysin digests **(Figure 1G-H)**. Although complete sequence coverage was obtained in some proteins with TrypLysC, inclusion of missed cleavages did not increase sequence coverage in several proteins. These analyses indicate Thermolysin may provide additional sequence coverage and proteomic depth unobtainable with TrypLysC.

**Figure 1.**
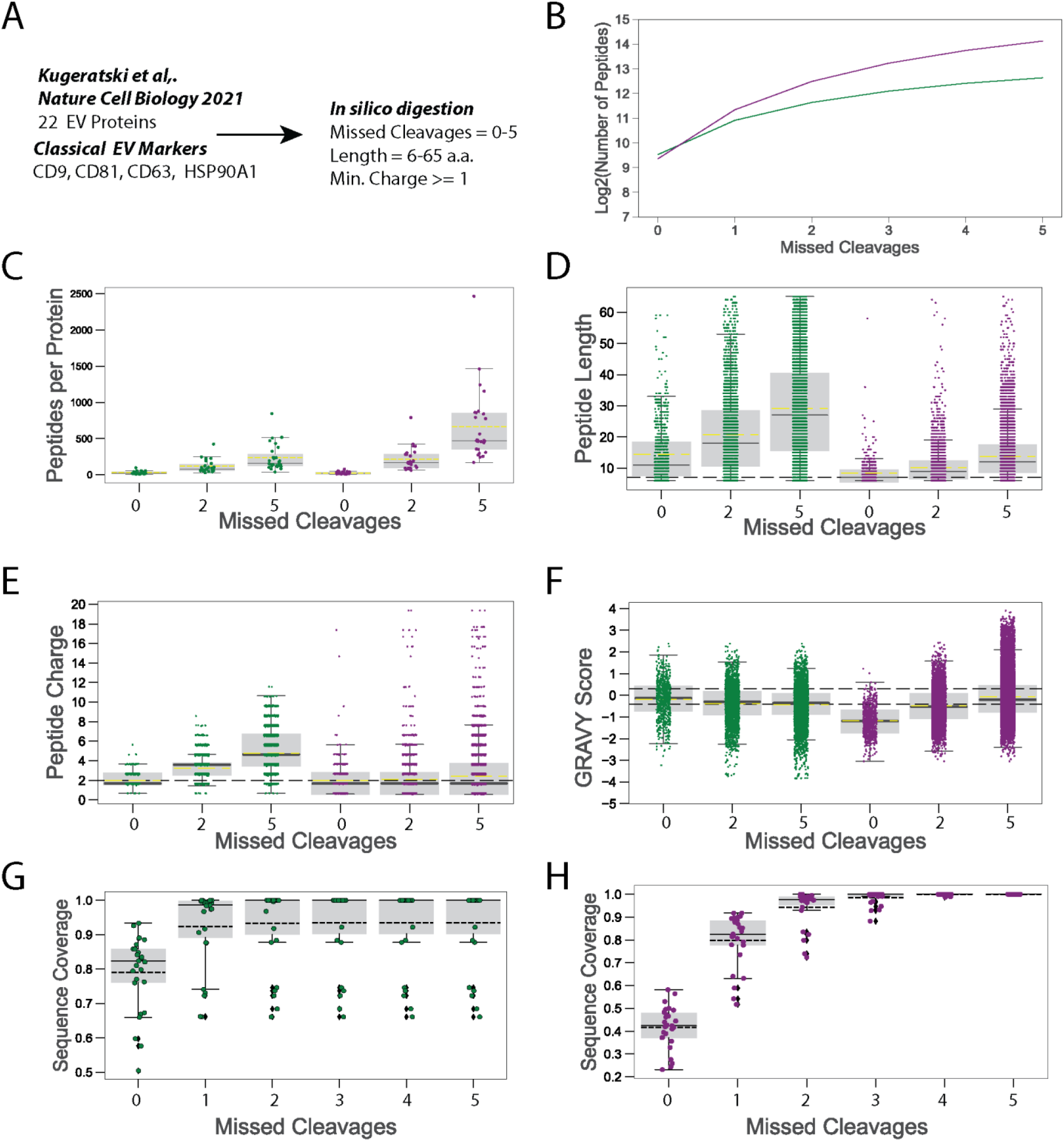
in silico proteolysis of core EV proteome. A) Workflow of in silico analysis using Kugeratski et al. at reference for “core” EV proteins. Relationship of missed cleavage sites and B) Number of unique peptides, C) peptide charge, and D) GRAVY score. Inclusion of missed cleavages improved sequence coverage of core EV proteins for G) TrypLysC and H) Thermolysin. Yellow dashed line indicates mean of data. TrypLysC = Green, Thermolysin = Purple.

### Thermolysin Proteolysis Enhances Proteomic Depth of Cell Line EVs

We sought to determine the experimental limitations of Thermolysin proteolysis by analyzing the proteomes of OC cell and ascites EVs. We hypothesized that Thermolysin would provide complementary sequence coverage to Trypsin/LysC and increase the number of identified proteins. To obtain missed cleavages sufficient for protein identification while avoiding the caveats of undigested proteins in nanoflow chromatography (i.e. blocked chromatography), we limited Thermolysin digestion to 1 hour at a 1:50 enzyme-to-protein ratio. Icelogo analysis of experimental peptides indicates that Thermolysin proteolysis was primarily restricted to the N-terminus of F/V/I/L, whereas cleavage at A/M was less commonly identified **(Figure 2A)**. Positional enrichment of amino acids determined that Thermolysin proteolysis was unlikely to occur when Proline was positioned at P-1 or P1 relative to the cleavage site (PO). This observation was similar to cleavage patterns of Tryp/LysC **(Figure 2B)**. Furthermore, cleavage site specificity of Thermolysin was positively associated with with Threonine (T) at positions −1 and 1 (P-1, P1), in addition to Aspartic Acid (D) at positions 2-4 (P2-4).

**Figure 2.**
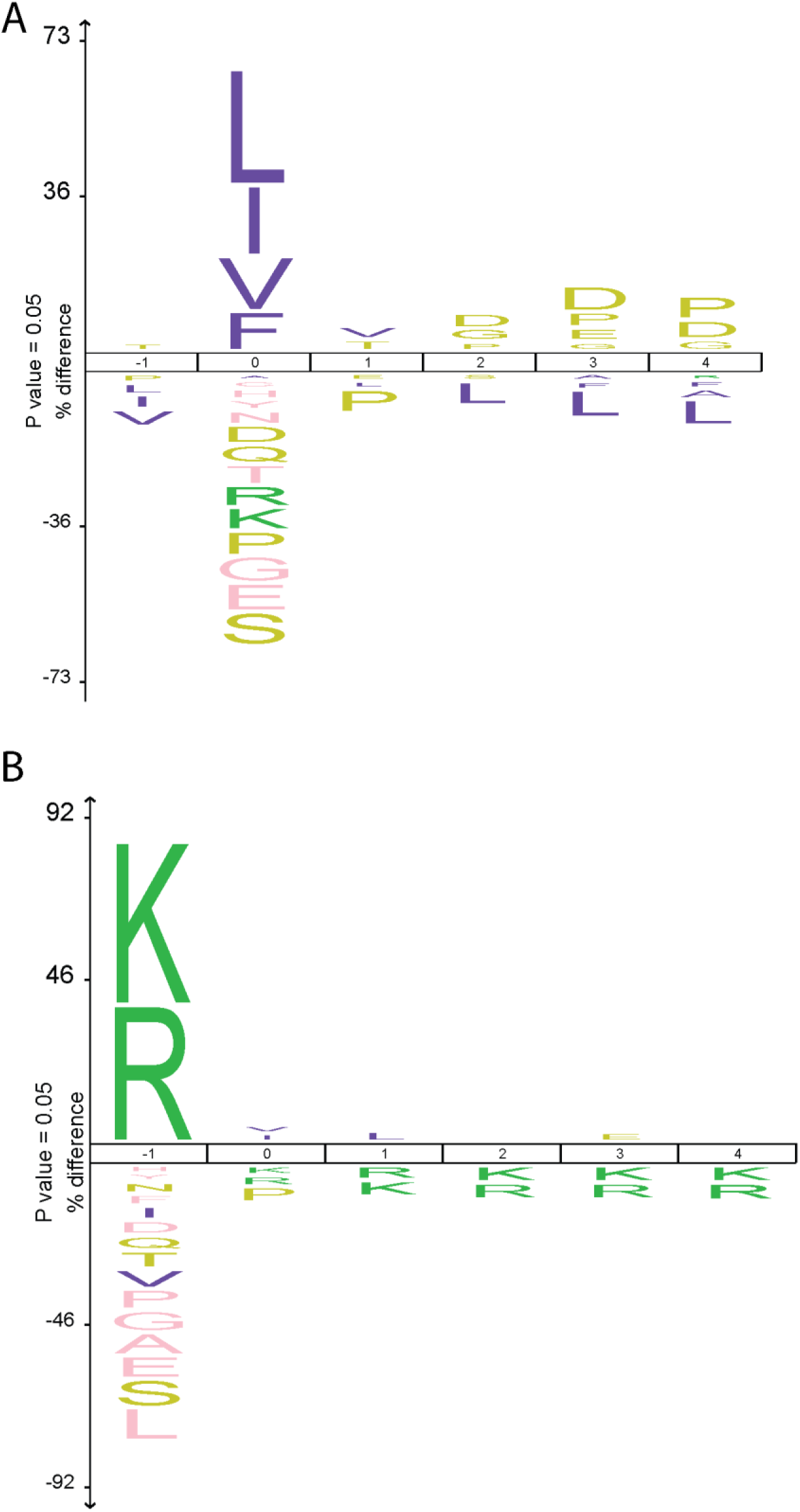
ICE logo analysis of cleavage sites. Cleavage site specificity for peptides generated by A) Thermolysin or B) TrypLysC identified by experimental LC-MS/MS.

We focused our initial assessments of Thermolysin-generated peptides using classical “shotgun” analyses with UPLC gradient lengths of 75 or 240 minutes **(Table 1; Supplemental Figure 1)**. Focusing on a 240-minute UPLC gradient, >3800 proteins were identified across OC cell line EVs. An average of 100 additional proteins were identified in Thermolysin digests, of which 84 of these proteins were identified in all OC cell lines. This included lymphocyte antigen 6E (Ly6E), a known driver of malignancy in breast and gynecological tumours^33, 34^. While fewer peptides and proteins were identified in Thermolysin compared to TrypLysC **(Figure 3A-B)**, the combination of both enzymes increased mean sequence coverage > 2-fold in both 75-minute and 240-minute gradients **(Figure 3C-D)**. GO Cellular Compartment (GOCC) enrichment analysis indicated that both Thermolysin and TrypLysC identified proteins involved with EV biogenesis (GO:0030126/ GO:0010008/GO:0030136), lipoproteins (GO:0034263), focal adhesions (GO:0005925), and cell-substrate interactions (GO:0030055) **(Figure 3E-F)**. Experimental proteolysis with Thermolysin or TrypLysC did not provide the sequence coverage of core EV proteins predicted with our in-silico analysis **(Figure 1G-H)**. Each enzyme was only able to provide partial sequence coverage regardless of cell line **(Figure 3G)**. These results indicate that TrypLysC and Thermolysin are complementary enzymes, albeit complete protein sequencing remains limited.

**Figure 3.**
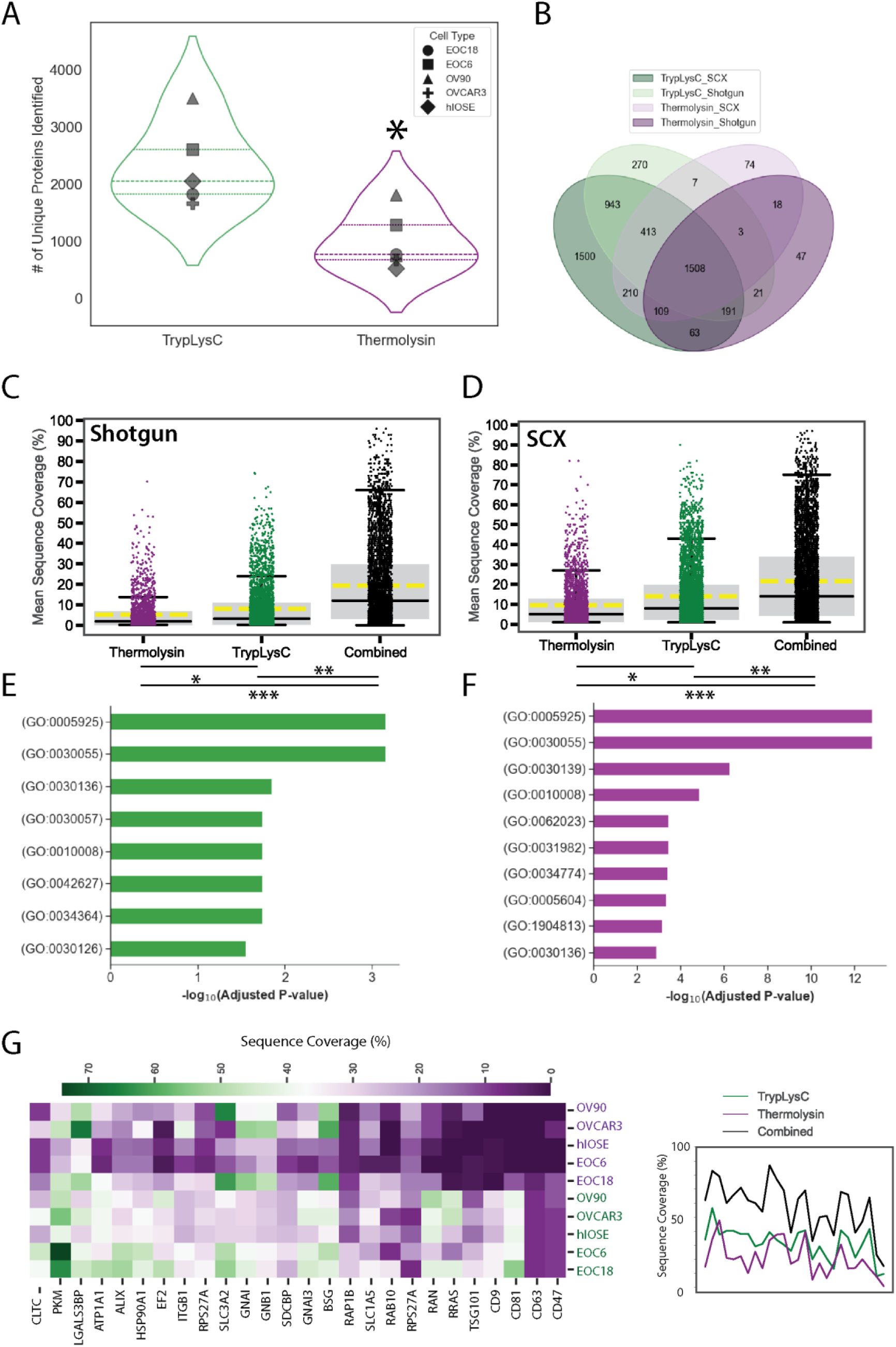
Thermolysin generates complementary peptides that increase proteomic depth and sequence coverage. EVs were isolated for OC cell lines using ultracentrifugation and analyzed by LC-MS/MS. A) Number of unique proteins identified in OC EVs was significantly less in Thermolysin digests. B) Venn diagram of proteins identified by shotgun or SCX analysis. C-D) TrypLysC provided increased sequence coverage compared to Thermolysin, regardless of MS/MS technique. Yellow dashed line indicates mean sequence coverage. E) GO Cellular Component analysis identified an enrichment of EV biogenic machinery related to EV biogenesis (GO:0030126/ GO:0010008/GO:0030136), lipoproteins (GO:0034263), focal adhesions (GO:0005925), and cell-substrate interactions (GO:0030055). G) Heatmap showing distinct sequence coverage profiles of core EV proteins between TrypLysC and Thermolysin. Combined peptide identification increased sequence coverage for all core EV proteins. Mann-Whitney U-test was used for A, whereas Kruskal-Wallis H Test with Dunn’s post hoc test was used for C-D. * = P<0.05, ** = P <0.01, *** = P<0.001.

**Table 1.** Summary of LC-MS/MS analysis of OC cell line EVs.

| Gradient Length (mins) | Enzyme | Peptide Dimensionality | MS1 Scans† | MS2 Scans† | Peptides† | Protein IDs† | Sequence Coverage † |
| --- | --- | --- | --- | --- | --- | --- | --- |
| 75 | TrypLysC | Shotgun | 7079.0 ± 493.8 | 26512.8 ± 2867.5 | 9271.4 ± 2975.4 | 1739 ± 316.0 | 7.10 ± 10.03 |
| 75 | Thermolysin | Shotgun | 8099.6 ± 543.9 | 21552.2 ± 2805.9 | 3218.6 ± 1533.6 | 745 ± 220.0 | 5.09 ± 7.69 |
| 75 | Combined | Shotgun | n/a | n/a | n/a | **3238** | **17.9 ± 19.29** |
| 75 | TrypLysC | SCX | 30382.4 ± 1533.6 | 95264.8 ± 15076.39 | 20870 ± 6249.86 | 2833 ± 447.03 | 12.64 ± 13.86 |
| 75 | Thermolysin | SCX | 33472.2 ± 2846.64 | 81145.6 ± 14050.95 | 7330.6 ± 3671.1 | 1345 ± 377.65 | 9.07 ± 10.54 |
| 75 | Combined | SCX | n/a | n/a | n/a | **4793** | **21.41 ± 21.86** |
| 260 | TrypLysC | Shotgun | 16704 ± 2771.9 | 130036.4 ± 15474.0 | 13349.2 ± 4490.90 | 2141.2 ± 387.15 | 8.01 ± 10.94 |
| 260 | Thermolysin | Shotgun | 20586 ± 3719.27 | 110886.4 ± 18416.7 | 4682.2 ± 2634.60 | 972.8 ± 286.57 | 5.27 ± 7.94 |
| 260 | Combined | Shotgun | n/a | n/a | n/a | **3829** | **19.30 ± 19.84** |
| 260 | TrypLysC | SCX | 3218.6 ± 1533.6 | 3218.6 ± 1533.6 | 9271.4 ± 2975.4 | 4980 ± 14.75 | 13.96 ± 14.75 |
| 260 | Thermolysin | SCX | 3218.6 ± 1533.6 | 3218.6 ± 1533.6 | 3218.6 ± 1533.6 | 1084 ± 460 | 9.48 ± 10.86 |
| 260 | Combined | SCX | n/a | n/a | n/a | **5088** | **21.66 ± 21.25** |

Abbreviations: SCX = Strong Cation Exchange , n/a = not applicable ; †Values presented as mean ± SD

### Offline Fractionation Using Strong Cation Exchange Increase Proteomic Depth

We hypothesized that decreasing peptide complexity would significantly improve peptide and protein identification within Thermolysin digests. Accordingly, we employed offline fractionation using SCX stagetips prepared into 200µl pipette tips **(Supplemental Figure 3)**. This method was chosen due to the theoretical diversity of charged species generated by Thermolysin **(Figure 1E)**. Relative to unfractionated peptide pools, SCX increased the identification of peptides across the whole gradient for both TrypLysC and Thermolysin **(Figure 4A)**. Peptides identified from Thermolysin digests typically contained 2-3 missed cleavage sites, albeit some peptides were identified with 4+ missed cleavages **(Figure 4B)**. These observations were in stark contract with TrypLysC, which primarily produced peptides with <2 missed cleavages. The addition of offline peptide fractionation led to the identification of >13,000 unique peptides in Thermolysin and >20,000 peptides in TrypLysC relative to shotgun analysis. In total, we identified >80,000 peptides, of which <0.3% were identified as contaminating or “carry over” peptides **(Figure 4C)**. Compared to shotgun analysis, the diversity of peptides obtained with SCX led to differences in the average mass (m/z) of peptides identified from TrypLysC or Thermolysin pools **(Figure 4D)**. SCX increased the charge distribution of peptides in Thermolysin pools, whereas this was not observed with TrypLysC. Differential m/z profiles between shotgun and SCX translated to peptides from Thermolysin that were statistically similar in length but were slightly more hydrophilic **(Figure 4F-G)**. In contrast, SCX of TrypLysC peptides were statistically smaller in length but also more hydrophilic than their shotgun counterparts. These observations do not align with our *in silico* analyses, which could be explained by poor ionization, fragmentation, or loss of peptides during sample preparation. In support of our hypothesis, decreasing peptide complexity using SCX increased the total number of unique proteins and average sequence coverage relative to unfractionated peptides **(Supplemental Figure 2A)**. Considering SCX requires 5x the instrument time for MS/MS acquisition versus shotgun analysis (6 versus 1 sample injections), the number of proteins identified per MS time was comparable between methodologies.

**Figure 4.**
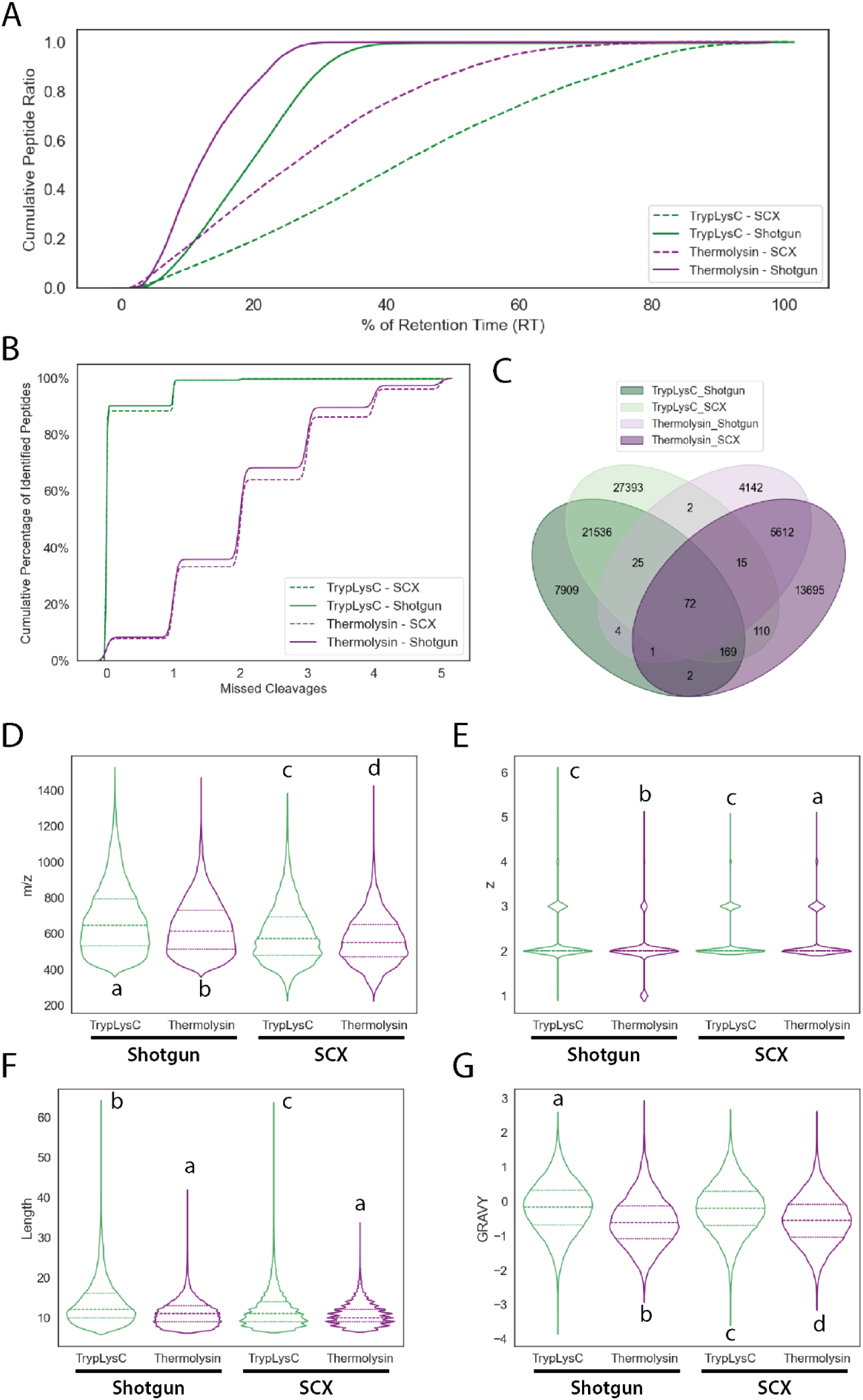
Strong cation exchange improves peptide identification and proteomic depth of OC EVs. SCX peptide fractionation was employed to reduce peptide complexity and increase proteomic depth in cell line EVs, reference *Table 1*. A) Relative retention times for peptides identified with or without SCX fraction. B). Relationship of missed cleavages to identification of unique peptides. C) Venn diagram showing overlap of proteins identified by SCX or shotgun analysis. Peptide characteristics such as D) m/z, E) charge, F) length, or G) GRAVY score. Data in D-G represented as violin plot with quartiles highlighted by horizontal lines. Annotated letters (a,b,c,d) indicate statistical grouping by Kruskal-Wallis H Test with Dunn’s post hoc test.

### Mining the Proteome of Primary Ascites Fluid EVs

To assess the applicability of Thermolysin proteolysis for biomarker discovery, we analyzed EVs isolated from clinical ascites fluid samples of women with HGSC. In contrast to cell line EVs, the total number of unique peptides was increased in Thermolysin samples relative to TrypLysC **(Figure 5A)**. Like cell line EVs, peptides generated by TrypLysC contained < 3 missed cleavages, whereas peptides with 4+ missed cleavages sites were confidently identified from Thermolysin digests with *de novo* sequencing **(Figure 5B)**. Peptides from Thermolysin digests typically eluted earlier in our chromatography relative to TrypLysC **(Figure 5C)**. These observations were consistent with the elution profile of Thermolysin peptides from SCX stagetips and coincided with the identification of more small hydrophilic peptides relative to TrypLysC. **(Figure 5D-F)**. Although we identified more peptides in Thermolysin digests, this did not translate to an increase in the number of unique proteins relative to TrypLysC **(Figure 5G)**. We speculate the discrepancy between elevated peptide numbers and proteins identification to be a result of the large dynamic range of proteins found in ascites EVs and the increased number of peptides per protein generated by Thermolysin with high missed cleavage rates **(Figure 1C)**. Together, peptide complexity and identification would be skewed to proteins at higher concentrations when digesting biofluids (e.g. plasma, ascites, etc.). Consistent with previous reports, identified peptides and proteins were primarily identified in the first few fractions of SCX fractions **(Figure 5H)**. Inclusion of proteins identified by Thermolysin increased to total proteomic depth by ∼19% compared to TrypLysC alone **(Figure 5I)**. GO Biological Process enrichment analysis of the 258 unique proteins identified by Thermolysin indicated involvement in preventing both endothelial and epithelial cell apoptosis **(Figure 5J)**. This included proteins such as IL-4 and ICAM1/CD54 that are previously identified in EVs^35, 36^ and elevated in the ascites of carcinoma vs benign disease^37^. GOCC analysis suggest the majority of these proteins to originate from lytic vacuoles or axonal projections **(Supplemental Figure 2B)**. We did not observe a bias of Thermolysin towards membrane proteins in Ascites EVs (**Supplemental Figure 2C)**, however we were able to increase the total number by ∼20% from Trypsin alone. Collectively, these results support our speculation that Thermolysin can provide additional insights into the proteomic landscape of EVs.

**Figure 5.**
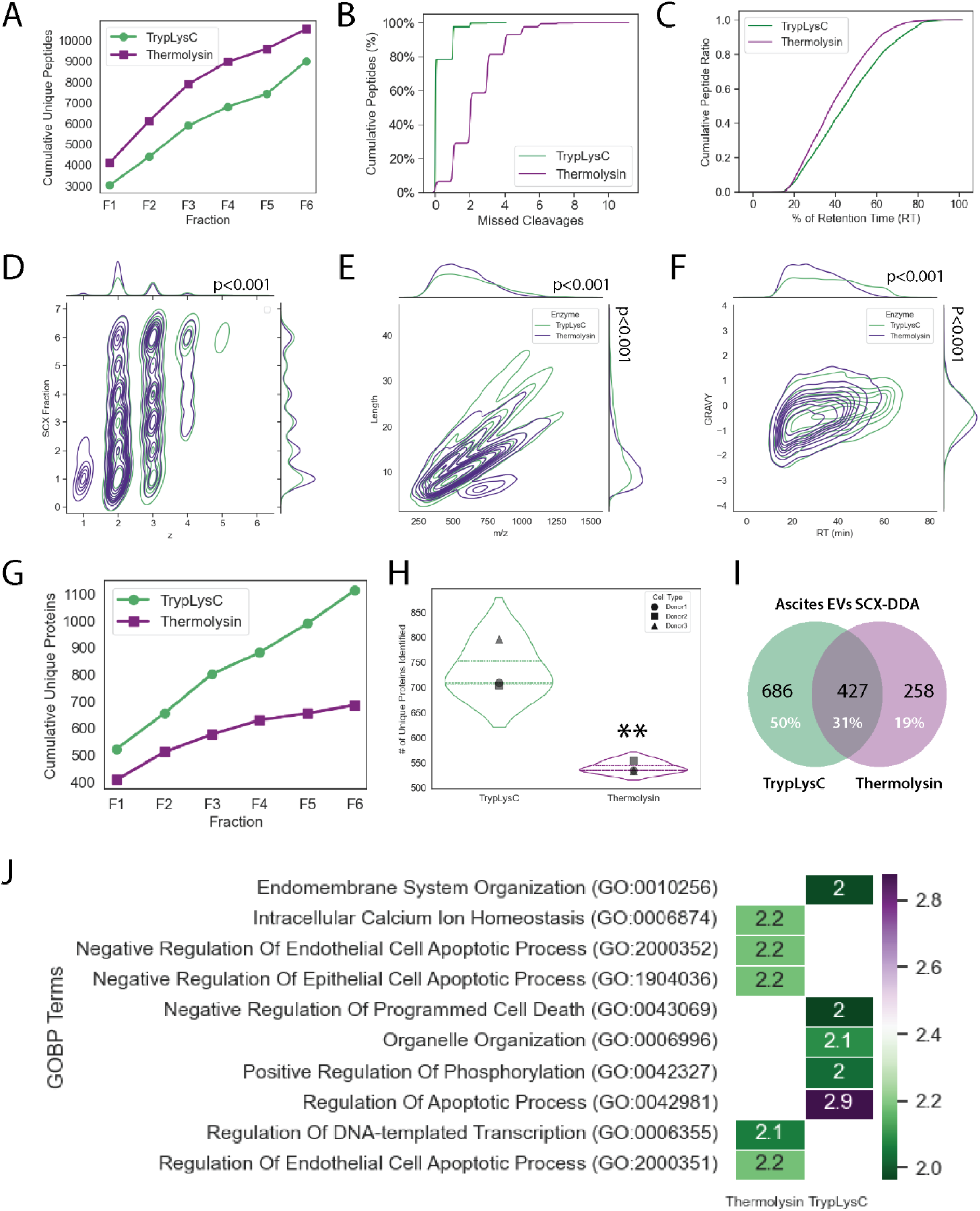
Mining the proteome of ascites EVs using SCX and DDA analysis. A) Total number of peptides identified in SCX fractions. B) Elution profiles. C) Relationship of peptide identification to number of missed cleavages. D) Relationship of peptide charge (z) and elution from SCX stagetips, E) peptide m/z vs length. F) chromatography and GRAVY scores. G) Relationship between protein unique protein identification and SCX fraction. H) Total number of unique proteins, I) Venn diagram showing overlap of proteins identified between TrypLysC and Thermolysin. J) GO Biological Process enrichment analysis for proteins exclusively identified by Thermolysin versus TrypLysC proteolysis. P-values in D-F indicate statistical analysis Mann-Whitney U test. ** = P <0.01

### Thermolysin proteolysis increases proteomic depth obtained with data-independent acquisition

We speculated that the rate and sensitivity of precursor selection in DDA mode might limit peptide/protein identification in Thermolysin samples. DIA is an alternative technique to DDA, where unbiased isolation and fragmentation of ions maximizes the speed at which MS/MS spectra are obtained^38, 39^. In theory, DIA should increase the detection of low-abundant peptides with similar biophysical properties to peptides derived from high-abundant proteins (i.e., albumin)^40, 41^ and also limit the number of missing values between samples^39^. For these reasons, we explored the use of SCX-DDA acquisitions to build a spectral library for wide-window DIA analyses using a 75-minute gradient. In contrast to DDA methods, we identified a comparable number of proteins from Thermolysin peptides relative to TrypLysC when samples were analyzed in DIA **(Figure 6A)**. Principal component analysis indicates that each enzyme produced a unique proteomic fingerprint for Ascites EVs separated on Principal Component 1, while differences between EV profiles of donors were retained as indicated by Principal Component 2 **(Figure 6C)**. Importantly, we demonstrate technical replicates injected following two separate instrument calibrations retained similar proteomic profiles regardless of proteolytic enzyme. In the context of proteome ‘mining’, Thermolysin provided 203 additional proteins that resulted in a ∼30% increase in proteomic depth obtained with DIA compared to TrypLysC alone **(Figure 6D)**. Despite distinct protein identification between the two enzymes, the distribution of sequence coverage obtained for TrypLysC was elevated (16.67 ± 15.68 vs 14.07 ± 15.68) compared to Thermolysin **(Figure 6D)**. These results indicate that Thermolysin is a useful proteolytic tool to complement TrypLysC for mining the proteome of biofluid EVs using DIA, albeit some limitations must be considered.

**Figure 6.**
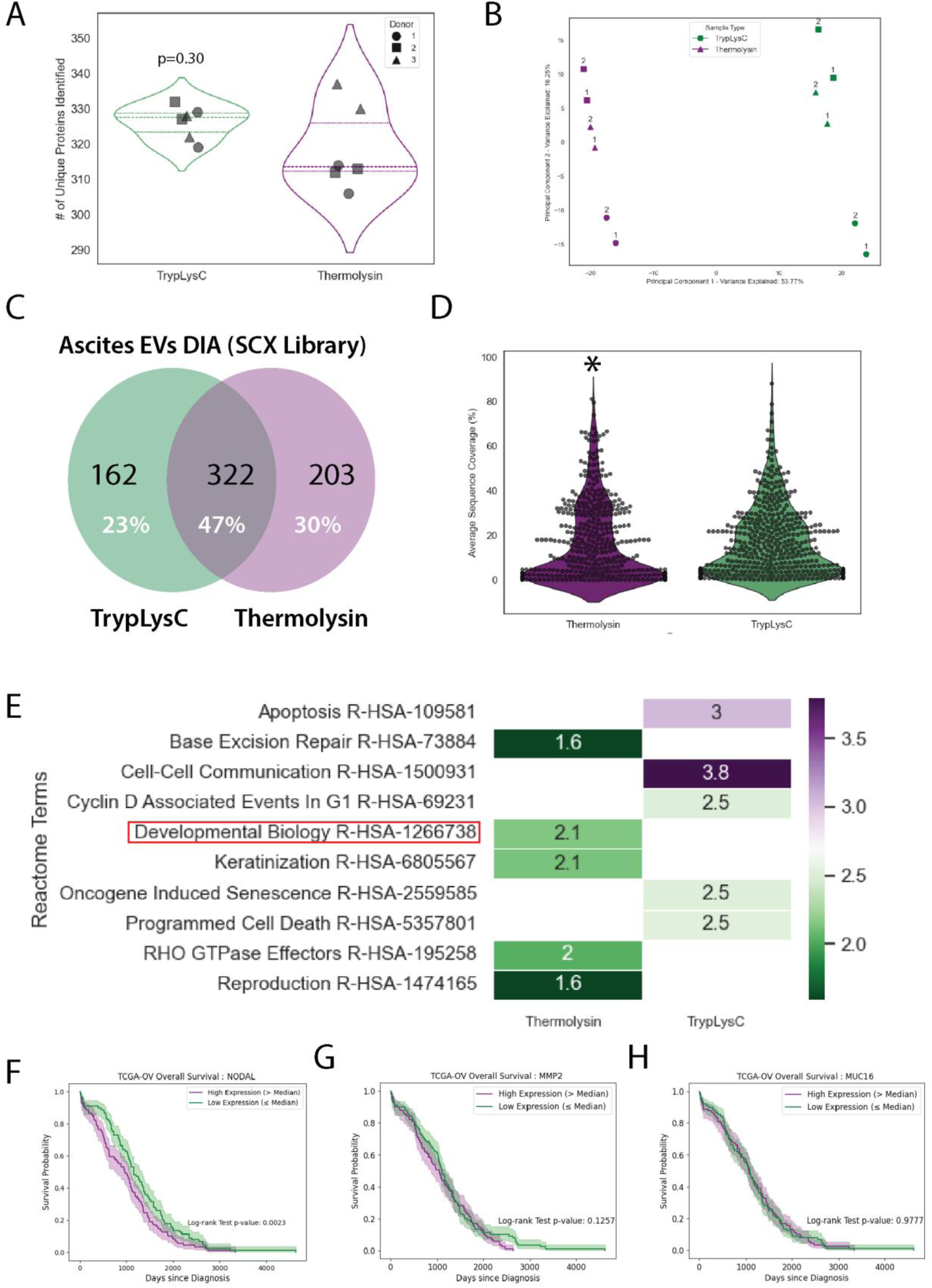
Data Independent Analysis of Ascites EVs. A) Total number of unique proteins identified by wide-window DIA. B) Principal component analysis of ascites EVs (n-3). Each donor and enzyme combination were analyzed as technical duplicates indicated as 1 and 2. C) Venn diagram of the number of unique proteins identified between enzymes. D) Sequence coverage of identified proteins using each enzyme. E) Reactome enrichment analysis of proteins uniquely identified by Thermolysin or TrypLysC proteolysis. Developmental Biology proteins were highlighted in Supplemental Figure 3. TCGA-OV analysis of F) NODAL, G) MMP2 and H) MUC16. P-values in D-F indicate statistical analysis Mann-Whitney U test. * = P <0.05

### Increased NODAL is Indicative of Poor Clinical Prognosis

We next sought to validate the biological significance of our proteomic findings in relation to OC progression, microenvironment, and recurrence. We focused our enrichment analysis on proteins uniquely identified by Thermolysin versus TrypLysC in our DIA analysis. Proteins associated with developmental biology were significantly enriched within Thermolysin digests **(Figure 6E)**, although several proteins were also identified in TrypLysC **(Supplemental Figure 3)**. Metalloproteinase 2 (MMP2) was exclusively detected in TrypLysC digests and projected externally from EV membranes in both healthy and pathological conditions to remodel the tissue microenvironment^42, 43^. NODAL, a member of the Transforming Growth Factor-Beta (TGF-β) superfamily, was exclusively detected in Thermolysin digests and is known to directly promote the progression of several cancers in response to hypoxia^44–46^. NODAL is pivotal in embryonic development and pluripotency; however, it re-emerges in adult pathologies to orchestrate growth-promoting programs. Its aberrant expression in cancers, including OC, is linked to aggressive metastasis and recurrence, highlighting its potential as a therapeutic target^46, 47^. To our knowledge, this is the first study to provide evidence that NODAL may be integrated into ascites EVs.

TCGA-OV analysis indicate that high NODAL expression is significantly associated with poor overall survival (OS). In contrast, MMP2 and current OC biomarker MUC16^48^ (CA125) did not provide this predictive power **(Figure 6F-G)**. To further understand our findings in the context of the ascites microenvironment, we quantified RNA from the AOCS primary ascites cohort using digital droplet PCR **(Figure 7A-C)**. NODAL RNA expression in ascites fluid was detected at low levels (114.2 ± 228.7, Mean ± SD) in agreement previously studies in breast and skin cancer that suggest NODAL expression is temporal and restricted to the local tumour microenvironment in adult tissues^44^. Regardless, we observed that increase NODAL mRNA in ascites fluid corresponds to poor OS (p=0.06) and progression-free survival (p=0.03), reaffirming that NODAL has an important role in tumour progression. These results were supported by the observation that NODAL mRNA was not significantly different between primary tumour and recurrence (p=0.62), was not induced with neoadjuvant chemotherapy (p=0.64) and did not deviate between alive and deceased patients (p=0.52) **(Figure 7D-E)**. Our current efforts are focused on elucidating the biological functions of NODAL in OC, specifically in relation to cellular plasticity and chemoresistance. Taken together, the results obtained within this study suggest a clinically relevant role of NODAL in the progression of OC.

**Figure 7.**
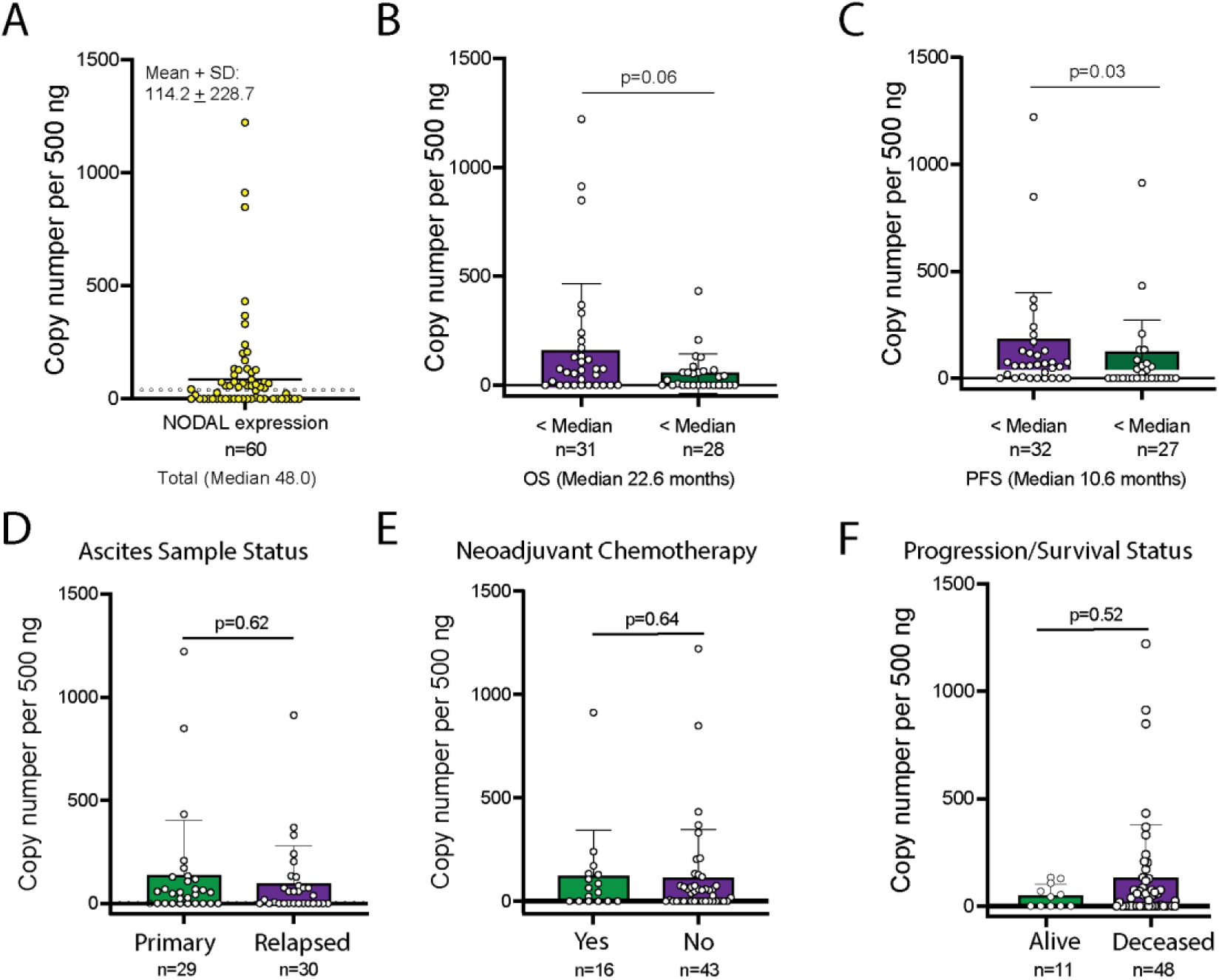
Elevated NODAL mRNA in ascites fluid corresponds to poor clinical outcomes in OC. NODAL mRNA was quantified in clinical ascites fluid (AOCS) using digital droplet PCR. A) NODAL mRNA copy number per 500ng of RNA. Increased NODAL identification was associated with B) overall survival and C) progression free survival. NODAL mRNA was not influenced by D) ascites status, E) chemotherapy, or survival status. P-values in D-F indicate statistical analysis Mann-Whitney U test.

## Discussion

The application of Thermolysin in the proteomic analysis of EVs from OC ascites fluid has yielded significant insights while also highlighting new challenges and opportunities. Thermolysin’s broad specificity for cleavage at the N-termini of hydrophobic residues (FMVAIL) contrasts sharply with the more restricted specificity of Trypsin/LysC, which cleaves at C-terminal of lysine (K) and arginine (R) residues. This broader specificity will result in a greater number of potential cleavage sites for Thermolysin that translates to an exponential increase in the complexity of peptide pools, particularly when missed cleavages are permitted. This heightened complexity will likely overwhelm current mass spectrometric capabilities, especially under data-dependent acquisition (DDA) settings, where the fixed acquisition speed and precursor selection may not limit unique peptide identification. To partition increased peptide complexity introduced by Thermolysin, we implemented SCX fractionation to segregate peptides by charge. Although SCX successfully reduced sample complexity and enhanced proteomic depth, alternative fractionation methods such as high-pH reverse phase chromatography might offer superior performance in decomplexifying peptide pools. Our study was limited by the absence of advanced mass spectrometric technologies such as FAIMS (Field Asymmetric Ion Mobility Spectrometry)^14^, ETD (Electron Transfer Dissociation)^49^, or state-of-the-art nanoflow chromatography that could further increase proteomic depth. The incorporation of these technologies could significantly enhance the resolution and sensitivity of our analyses, potentially improving the quantification and identification of peptides generated by Thermolysin. Mining the proteome of complex samples with complementary enzymes presents a valuable opportunity to explore the impact of increased sequence coverage on the identification of novel post-translational modifications (PTMs)^50^. While our current investigation did not delve deeply into PTMs, the enhanced sequence coverage provided by Thermolysin suggests a promising avenue for future research. Identifying and characterizing novel PTMs within the EV proteome could provide crucial insights into the molecular mechanisms of OC and uncover new therapeutic targets. Additionally, the method of EV isolation significantly influences the outcome of proteomic analyses^26^. While our study employed standard EV isolation techniques, exploring alternative methods could further improve the efficiency and specificity of proteomic profiling. Techniques such as size-exclusion chromatography or immunoaffinity capture, which offer higher purity of EVs would likely provide additional proteomic cargo not identified in EVs isolated by UC. The integration of alternative technologies at all levels of analysis would increase the prospective pool of clinically relevant biomarkers and therapeutic targets that remain elusive with current EV isolation or MS/MS techniques.

In conclusion, while the use of Thermolysin has revealed new dimensions in the proteome of OC EVs, it also underscores the need for continuous technological advancements and methodological refinements in proteomics. Future studies equipped with superior peptide fractionation techniques, multidimensional mass spectrometry analyses, and enhanced EV isolation methods are poised to further unravel the complex proteome of OC EVs. This study highlighted the application of complementary proteolysis to mine the proteome of EVs isolated from cell culture and clinical samples. We exemplified the importance of these findings by validating NODAL as a clinically relevant protein using TGCA and AOCS cohorts. These advancements will pave the way for groundbreaking discoveries in OC diagnostics and therapy, ultimately improving patient outcomes for a currently challenging disease.

## Acknowledgements

The Australian Ovarian Cancer Study gratefully acknowledges additional support from Ovarian Cancer Australia and the Peter MacCallum Foundation. The AOCS also acknowledges the cooperation of the participating institutions in Australia and acknowledges the contribution of the study nurses, research assistants and all clinical and scientific collaborators to the study. The complete AOCS Study Group can be found at https://smex-ctp.trendmicro.com:443/wis/clicktime/v1/query?url=www.aocstudy.org&umid=83d7b2f5-ae02-459b-a168-e82a948f529f&auth=6ee66d642a212b82964c9073f0dd934b55317413-a35557dabe6675b967bae2754a69d75c0a5252b5. We would like to thank all of the women who participated in these research programs.

## References

1. Dupree EJ, Jayathirtha M, Yorkey H, Mihasan M, Petre BA and Darie CCJP. A critical review of bottom-up proteomics: the good, the bad, and the future of this field. 2020;8:14.

2. Leutert M, Entwisle SW, Villén JJM and Proteomics C. Decoding post-translational modification crosstalk with proteomics. 2021;20.

3. Manuel JM, Guilloy N, Khatir I, Roucou X and Laurent BJFiG. Re-evaluating the impact of alternative RNA splicing on proteomic diversity. 2023;14:1089053.

4. Canty EG and Kadler KEJJocs. Procollagen trafficking, processing and fibrillogenesis. 2005;118:1341–1353.

5. Hu Q, Noll RJ, Li H, Makarov A, Hardman M and Graham Cooks RJJoms. The Orbitrap: a new mass spectrometer. 2005;40:430–443.

6. Makarov A, Denisov E, Kholomeev A, Balschun W, Lange O, Strupat K and Horning SJAc. Performance evaluation of a hybrid linear ion trap/orbitrap mass spectrometer. 2006;78:2113–2120.

7. Sjödin MO, Wetterhall M, Kultima K and Artemenko KJJocB. Comparative study of label and label-free techniques using shotgun proteomics for relative protein quantification. 2013;928:83–92.

8. Timp W and Timp GJSA. Beyond mass spectrometry, the next step in proteomics. 2020;6:eaax8978.

9. Wright BW, Yi Z, Weissman JS and Chen JJTiCB. The dark proteome: translation from noncanonical open reading frames. 2022;32:243–258.

10. Kuljanin M, Dieters-Castator DZ, Hess DA, Postovit LM and Lajoie GAJP. Comparison of sample preparation techniques for large-scale proteomics. 2017;17:1600337.

11. Berberich MJ, Paulo JA and Everley RAJJopr. MS3-IDQ: Utilizing MS3 spectra beyond quantification yields increased coverage of the phosphoproteome in isobaric tag experiments. 2018;17:1741–1747.

12. Navarrete-Perea J, Yu Q, Gygi SP and Paulo JAJJopr. Streamlined tandem mass tag (SL-TMT) protocol: an efficient strategy for quantitative (phospho) proteome profiling using tandem mass tag-synchronous precursor selection-MS3. 2018;17:2226–2236.

13. Park J, Yu F, Fulcher JM, Williams SM, Engbrecht K, Moore RJ, Clair GC, Petyuk V, Nesvizhskii AI and Zhu YJAC. Evaluating linear ion trap for MS3-based multiplexed single-cell proteomics. 2023;95:1888–1898.

14. Kaulich PT, Cassidy L, Winkels K and Tholey AJAC. Improved identification of proteoforms in top-down proteomics using FAIMS with internal CV stepping. 2022;94:3600–3607.

15. Pfammatter S, Bonneil E, McManus FP, Prasad S, Bailey DJ, Belford M, Dunyach J-J, Thibault PJM and Proteomics C. A novel differential ion mobility device expands the depth of proteome coverage and the sensitivity of multiplex proteomic measurements. 2018;17:2051–2067.

16. Cristobal A, Marino F, Post H, van den Toorn HW, Mohammed S and Heck AJJAc. Toward an optimized workflow for middle-down proteomics. 2017;89:3318–3325.

17. Tsiatsiani L and Heck AJJTFj. Proteomics beyond trypsin. 2015;282:2612–2626.

18. Liu Y, Engelman DM and Gerstein MJGb. Genomic analysis of membrane protein families: abundance and conserved motifs. 2002;3:1–12.

19. Bendz M, Skwark M, Nilsson D, Granholm V, Cristobal S, Käll L and Elofsson AJP. Membrane protein shaving with thermolysin can be used to evaluate topology predictors. 2013;13:1467–1480.

20. Saveliev S, Engel L, Strauss E, Jones R and Rosenblatt M. The Advantages to Using Arg-C, Elastase, Thermolysin and Pepsin for Protein Analysis. 2007.

21. Jevon D, Moon K-M, Foster LJ and Johnson JDJb. Identification of unusually disulphide-bonded insulin forms using mass spectrometry and thermolysin cleavage. 2022:2022.03.11.484026.

22. Rocha-Martin J, Fernández-Lorente G, Guisan JMJB and Biotransformation. Sequential hydrolysis of commercial casein hydrolysate by immobilized trypsin and thermolysin to produce bioactive phosphopeptides. 2018;36:159–171.

23. Jiang X, Yeung D, Liu Y, Spicer V, Afshari H, Lao Y, Lin F, Krokhin O and Zahedi RPJJoPR. Accelerating Proteomics Using Broad Specificity Proteases. 2024;23:1360–1369.

24. Welsh JA, Goberdhan DC, O’Driscoll L, Buzas EI, Blenkiron C, Bussolati B, Cai H, Di Vizio D, Driedonks TA and Erdbrügger UJJoev. Minimal information for studies of extracellular vesicles (MISEV2023): From basic to advanced approaches. 2024;13:e12404.

25. Yáñez-Mó M, Siljander PR-M, Andreu Z, Bedina Zavec A, Borràs FE, Buzas EI, Buzas K, Casal E, Cappello F and Carvalho JJJoev. Biological properties of extracellular vesicles and their physiological functions. 2015;4:27066.

26. Cooper TT, Dieters-Castator DZ, Liu J, Siegers GM, Pink D, Veliz L, Lewis JD, Lagugné-Labarthet F, Fu Y and Steed HJJoOR. Targeted proteomics of plasma extracellular vesicles uncovers MUC1 as combinatorial biomarker for the early detection of high-grade serous ovarian cancer. 2024;17:149.

27. Herrmann IK, Wood MJA and Fuhrmann GJNn. Extracellular vesicles as a next-generation drug delivery platform. 2021;16:748–759.

28. Cooper TT, Sherman SE, Bell GI, Dayarathna T, McRae DM, Ma J, Lagugné-Labarthet F, Pasternak SH, Lajoie GA, Hess DAJSC and Development. Ultrafiltration and injection of islet regenerative stimuli secreted by pancreatic mesenchymal stromal cells. 2021;30:247–264.

29. Ćulum NM, Cooper TT, Bell GI, Hess DA, Lagugné-Labarthet FJA and chemistry b. Characterization of extracellular vesicles derived from mesenchymal stromal cells by surface-enhanced Raman spectroscopy. 2021;413:5013–5024.

30. Ćulum NM, Cooper TT, Lajoie GA, Dayarathna T, Pasternak SH, Liu J, Fu Y, Postovit L-M and Lagugné-Labarthet FJA. Characterization of ovarian cancer-derived extracellular vesicles by surface-enhanced Raman spectroscopy. 2021;146:7194–7206.

31. Veliz L, Cooper TT, Grenier-Pleau I, Abraham SA, Gomes J, Pasternak SH, Dauber B, Postovit LM, Lajoie GA and Lagugné-Labarthet FJAs. Tandem SERS and MS/MS Profiling of Plasma Extracellular Vesicles for Early Ovarian Cancer Biomarker Discovery. 2024;9:272–282.

32. Kugeratski FG, Hodge K, Lilla S, McAndrews KM, Zhou X, Hwang RF, Zanivan S and Kalluri RJNcb. Quantitative proteomics identifies the core proteome of exosomes with syntenin-1 as the highest abundant protein and a putative universal biomarker. 2021;23:631–641.

33. Luo L, McGarvey P, Madhavan S, Kumar R, Gusev Y and Upadhyay GJO. Distinct lymphocyte antigens 6 (Ly6) family members Ly6D, Ly6E, Ly6K and Ly6H drive tumorigenesis and clinical outcome. 2016;7:11165.

34. Yeom CJ, Zeng L, Goto Y, Morinibu A, Zhu Y, Shinomiya K, Kobayashi M, Itasaka S, Yoshimura M and Hur C-GJO. LY6E: a conductor of malignant tumor growth through modulation of the PTEN/PI3K/Akt/HIF-1 axis. 2016;7:65837.

35. Casella G, Colombo F, Finardi A, Descamps H, Ill-Raga G, Spinelli A, Podini P, Bastoni M, Martino G and Muzio LJMT. Extracellular vesicles containing IL-4 modulate neuroinflammation in a mouse model of multiple sclerosis. 2018;26:2107–2118.

36. Wang L, Du D-D, Zheng Z-X, Shang P-F, Yang X-X, Sun C, Wang X-Y, Tang Y-J and Guo X-LJFiP. Circulating galectin-3 promotes tumor-endothelium-adhesion by upregulating ICAM-1 in endothelium-derived extracellular vesicles. 2022;13:979474.

37. Atta S, Kamel M, Mansour W, Hussein T, Maher K and Elrefaiy MAJDD. Ascitic fluid cytokines in chronic liver disease: a possible prognostic tool. 2021;39:534–539.

38. Gillet LC, Navarro P, Tate S, Röst H, Selevsek N, Reiter L, Bonner R, Aebersold RJM and Proteomics C. Targeted data extraction of the MS/MS spectra generated by data-independent acquisition: a new concept for consistent and accurate proteome analysis. 2012;11.

39. Krasny L and Huang PHJMo. Data-independent acquisition mass spectrometry (DIA-MS) for proteomic applications in oncology. 2021;17:29–42.

40. Pino LK, Just SC, MacCoss MJ, Searle BCJM and Proteomics C. Acquiring and analyzing data independent acquisition proteomics experiments without spectrum libraries. 2020;19:1088–1103.

41. Li J, He X, Deng Y and Yang CJM. An update on isolation methods for proteomic studies of extracellular vesicles in biofluids. 2019;24:3516.

42. Tang H, He Y, Li L, Mao W, Chen X, Ni H, Dong Y and Lyu FJEcr. Exosomal MMP2 derived from mature osteoblasts promotes angiogenesis of endothelial cells via VEGF/Erk1/2 signaling pathway. 2019;383:111541.

43. Shimoda M and Khokha RJBeBA-MCR. Metalloproteinases in extracellular vesicles. 2017;1864:1989–2000.

44. Quail DF, Siegers GM, Jewer M, Postovit L-MJTijob and biology c. Nodal signalling in embryogenesis and tumourigenesis. 2013;45:885–898.

45. Jewer M, Lee L, Leibovitch M, Zhang G, Liu J, Findlay SD, Vincent KM, Tandoc K, Dieters-Castator D and Quail DFJNc. Translational control of breast cancer plasticity. 2020;11:2498.

46. Quail DF, Taylor MJ, Walsh LA, Dieters-Castator D, Das P, Jewer M, Zhang G and Postovit L-MJMbotc. Low oxygen levels induce the expression of the embryonic morphogen Nodal. 2011;22:4809–4821.

47. Dieters-Castator D, Dantonio PM, Piaseczny M, Zhang G, Liu J, Kuljanin M, Sherman S, Jewer M, Quesnel K and Kang E-Y. Embryonic protein NODAL as a potential modulator of the tumour microenvironment: Breast cancer secretome reprogramming and fibroblast activation. CANCER RESEARCH. 2021;81.

48. Charkhchi P, Cybulski C, Gronwald J, Wong FO, Narod SA and Akbari MRJC. CA125 and ovarian cancer: a comprehensive review. 2020;12:3730.

49. Tu C, Li J, Young R, Page BJ, Engler F, Halfon MS, Canty Jr JM and Qu JJAc. Combinatorial peptide ligand library treatment followed by a dual-enzyme, dual-activation approach on a nanoflow liquid chromatography/orbitrap/electron transfer dissociation system for comprehensive analysis of swine plasma proteome. 2011;83:4802–4813.

50. Andaluz Aguilar H, Iliuk AB, Chen I-H and Tao WAJNp. Sequential phosphoproteomics and N-glycoproteomics of plasma-derived extracellular vesicles. 2020;15:161–180.

