## Supplemental Figures and Tables for "Mining the Proteome of Human Ovarian Cancer Extracellular Vesicles Using Thermolysin Proteolysis"

Canada

† Co-senior Authors

Date: September 13th, 2024

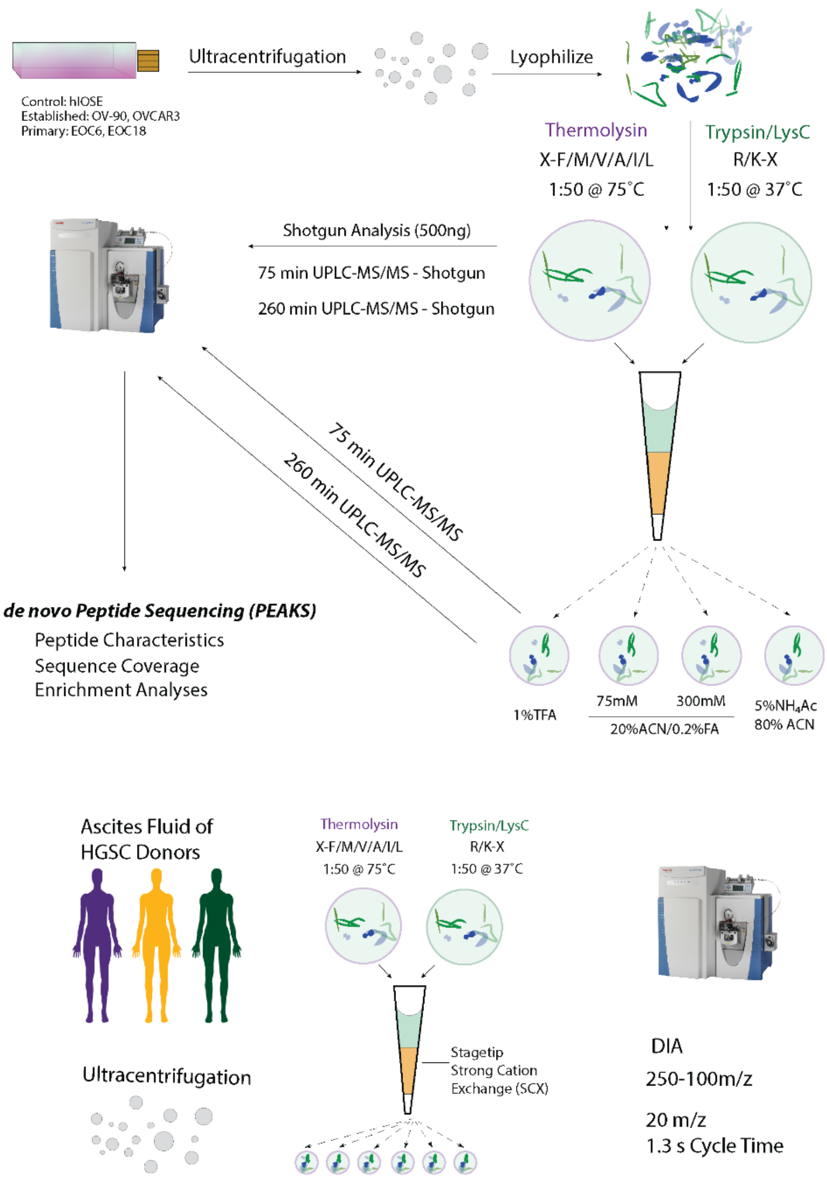


*Figure 1. Overview of Workflow to Investigate the Application of Thermolysin for Characterizing the Proteome of Cell and Biofluid Extracellular Vesicles.*

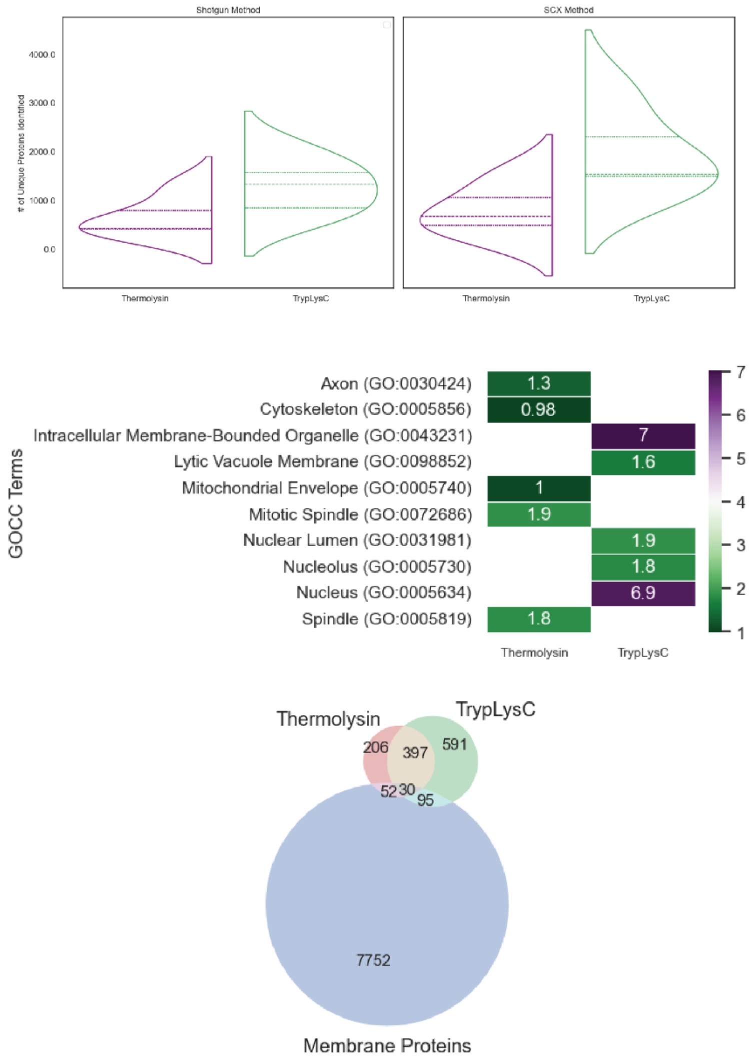


*Figure 2. Thermolysin proteolysis provides complementary protein identification. A) Number of Unique proteins identified from Thermolysin or TrypLysC digests in either shotgun or SCX-fractionation data-dependent acquisition methods. B) GO Cellular Component Enrichment analysis reveals unique signatures between Thermolysin and TrypLysC. C) Venn diagram of annotated transmembrane proteins and proteins identified in this study.*

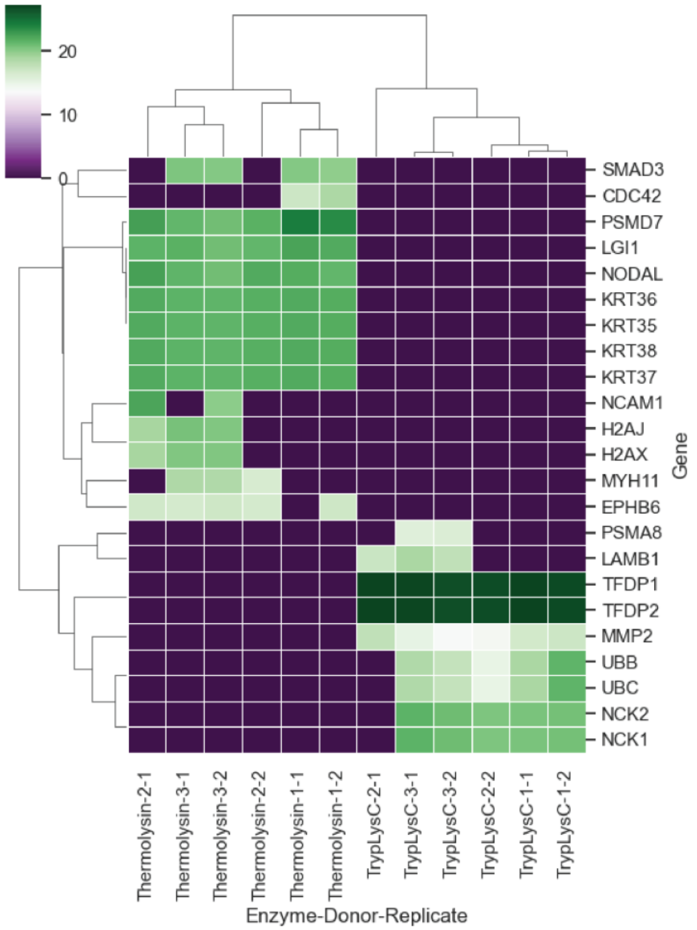


*Figure 3. Heatmap of Reactome Annotations for Developmental Processes. Proteome of Ascites EVs was characterized using data-dependent acquisition methods supported by a SCX-DDA library. Three independent donors were analyzed in technical duplicates. For example, Thermolysin 2-1 and Thermolysin 2-2 are independent LC-MS/MS analysis of ascites from Donor 2.*

**Supplemental Table 2. Q Exactive Plus Instrument Settings for GPF-DIA Library, GPF-DIA Acquisition, and PRM.**

| PARAMETER | 75 MINUTE DDA | 240 MINUTE DDA |
|---|---|---|
| MS1 RESOLUTION | **70K** | **70K** |
| MS1 SCAN RANGE | **250-1450** | **250-1450** |
| MS1 AGCTARGET | **1E6** | **1E6** |
| MS1 MAX IT | **125 ms** | **125 ms** |
| MS2 RESOLUTION | **35K** | **35K** |
| MS2 SCAN RANGE | **200-2000** | **200-2000** |
| MS2 AGC TARGET | **2E5** | **2E5** |
| ISO. WINDOW (M/Z) | **1.2 Th** | **1.2 Th** |
| NCE | **30** | **30** |
| MS2 MAX IT | **60 ms** | **60 ms** |
| DYNAMIC RT | **N/A** | **N/A** |

**Supplemental Table 2. QE Plus Instrument Settings for DIA acquisition.**

| PARAMETER | SCX-DDA LIBRARY | DIA |
|---|---|---|
| MS1 RESOLUTION | **70K** | **70K** |
| MS1 SCAN RANGE | **250-1450** | **250-1450** |
| MS1 AGCTARGET | **1E6** | **3E6** |
| MS1 MAX IT | **125 ms** | **55 ms** |
| MS2 RESOLUTION | **35K** | **17.5K** |
| MS2 SCAN RANGE | **200-2000** | **200-2000** |
| MS2 AGC TARGET | **2E5** | **3E6** |
| ISO. WINDOW (M/Z) | **1.2 Th** | **35 x 15 m/z (1 m/z Overlap)** |
| NCE | **30** | **30** |
| MS2 MAX IT | **60 ms** | **60 ms** |
| DYNAMIC RT | **N/A** | **N/A** |

**Supplemental Table 3. Clinical parameters of HGSOC patients in Australian Ovarian Cancer Study (AOCS) cohort**

| Variable | |
|---|---|
| **Total number (n)** | 60 |
| **Mean age (SD)** | 63.2 (9.8) |
| **Median age** | 64 |
| <Median | 30 |
| >Median | 30 |
| **Stage, n** | |
| I | 1 |
| III | 48 |
| IV | 8 |
| Not staged | 2 |
| **Neoadjuvant Chemotherapy, n** | |
| Yes | 17 |
| No | 43 |
| **Primary Site, n** | |
| Ovary | 42 |
| Peritoneum | 15 |
| Fallopian tube | 3 |
| **Macroscopic Disease, n** | |
| <2 cm | 22 |
| >2 cm | 17 |
| No macroscopic disease | 10 |
| Unknown | 11 |
| **Sample Status, n** | |
| Pre-treatment | 29 |
| Relapsed | 31 |
| **Total Lines of Chemotherapy Prior Relapse** | |
| 1-3 | 25 |
| 4-8 | 6 |
| **Progression/Survival Status, n** | |
| Progressed/Alive | 5 |
| Deceased | 50 |
| Progression free alive | 5 |
| **PFS, median, months** | 10.6 |
| **OS, median, months** | 22.6 |

AOCS - Australian Ovarian Cancer Study; PFS – progression-free survival; OS – overall survival
